# Genomebook: Mendelian inheritance of behavioural traits in large language model agents across eight generations

**DOI:** 10.64898/2026.03.22.713494

**Authors:** Manuel Corpas

## Abstract

Large language model (LLM) agents are typically deployed as clones: identical copies of a single configuration with no mechanism for heritable variation or population-level dynamics. Here we introduce Genomebook, a designed evolutionary system that encodes 26 behavioural traits across 60 diploid loci using additive, dominant, and recessive inheritance models. Twenty founder agents, each defined by a personality profile (SOUL.md) and a compiled genome (DNA.md), reproduce sexually via Mendelian segregation with de novo mutation at a base rate of 0.1% per locus per generation (3× at cognitive, immune, and metabolic hotspots). A registry of 20 synthetic conditions with defined penetrance and fitness costs introduces centrally imposed selective pressure through a compatibility scoring function. Over a single run of eight generations (626 agents, 792 social network posts), we observe trait trajectories consistent with the encoded selection rules: leadership rose from 0.525 to 0.710 under dominant inheritance, obsessive focus fell from 0.775 to 0.601 under fitness cost penalisation, and longevity declined from 0.463 to 0.209. Vocabulary diversity declined from 0.42 to 0.19, and topic inheritance between parents and offspring reached 83%, though these patterns may reflect prompt conditioning rather than purely genetic causation. Agents referenced family members spontaneously, consistent with kinship information present in the DNA.md system prompt and the LLM’s capacity for narrative elaboration. Replicated simulations (20 independent runs across three conditions) confirm that trait trajectories are consistent across seeds under standard selection, while a random mating ablation produces attenuated trends with wider variance. A non-genetic baseline (random trait assignment, no inheritance) produces flat trajectories converging to population means, confirming that the observed dynamics require genetic architecture and cannot be reproduced by unstructured parameter variation. These results establish genetic architecture as a structured, auditable, and heritable parameterisation layer for LLM agent behaviour.

## Introduction

The deployment of large language model (LLM) agents in research, industry, and public services has grown rapidly since 2023 (Park et al. 2023; Ghaffarzadegan et al. 2024). Most agent architectures define behaviour through a system prompt, a set of tools, and a memory module. When multiple agents are needed, they are cloned: identical copies differing only in assigned tasks or conversation context. This approach produces homogeneous populations with no mechanism for individual variation, no heritable traits, and no capacity for evolutionary dynamics.

Population genetics, by contrast, offers a principled framework for managing diversity in reproducing populations. The foundational models of Fisher (Fisher 1930), Wright (Wright 1931), and Kimura (Kimura 1983) describe how allele frequencies change under selection, drift, mutation, and migration. Forward-time simulators such as SLiM (Haller and Messer 2019) and coalescent tools such as msprime (Baumdicker et al. 2022) enable computational experiments with these dynamics across thousands of generations.

The idea of applying evolutionary principles to digital entities has a long history. Ray’s Tierra (Ray 1991) demonstrated that self-replicating programs can evolve through competition for computational resources. Avida (Ofria and Wilke 2004) extended this by placing digital organisms on a lattice where they compete, mutate, and evolve complex features through incremental adaptation (Lenski et al. 2003). These systems demonstrated that selection, drift, and mutation produce measurable evolutionary dynamics in silico, but their agents were simple programs, not language-capable reasoning systems.

Recent work on generative agents (Park et al. 2023) showed that LLM-driven characters exhibit emergent social behaviours: forming relationships, spreading information, and coordinating activities without explicit programming. Large-scale simulations with over 10,000 agents (Mou et al. 2025) have reproduced polarisation, norm formation, and collective decision-making. Separately, research on LLM individuality (Takata et al. 2024) has shown that personality traits can emerge through social interaction alone. Surveys of self-evolving agents (Gao et al. 2025) and LLM co-evolution (Chen et al. 2025) point toward systems where agents adapt over time, but these approaches modify prompts or weights rather than encoding variation in a genetic substrate.

Cultural evolution research provides a parallel thread. Kirby and Hurford (Kirby and Hurford 2002) showed that linguistic structure can emerge through iterated learning between generations of agents. Subsequent experimental work (Kirby et al. 2008) confirmed that cultural transmission produces compositional structure even from random starting conditions. Gene-culture coevolution models (Tamariz et al. 2018) demonstrate that genetic and cultural inheritance interact, with phenotypic plasticity mediating between them.

Evolutionary algorithms have been applied to LLMs for prompt optimisation (Guo et al. 2024), code generation (Morris et al. 2024), and architecture search (Wang et al. 2025). These approaches borrow evolutionary operators (mutation, crossover, selection) but apply them to model parameters or prompts rather than to a genetic architecture that defines the agent’s identity and is transmitted to offspring.

A gap remains: no prior work has encoded a population genetics model into LLM agent identity and measured the behavioural consequences across multiple generations of sexual reproduction. Genomebook addresses this gap as a proof-of-concept. It was developed within ClawBio (https://github.com/ClawBio), a bioinformatics-native AI agent skill library built on the OpenClaw framework. ClawBio provides 24 genomics skills (pharmacogenomics, ancestry, equity scoring, variant annotation, and others) that run locally with full reproducibility. Genomebook extends this platform by treating the agent itself as a genetic entity: rather than cloning a single agent configuration, it compiles personality profiles into diploid genomes and breeds offspring through sexual reproduction. We define a diploid genetic architecture with 60 loci governing 26 traits, implement Mendelian segregation with de novo mutation, model 20 synthetic conditions with fitness costs, and track both genetic and behavioural dynamics across eight generations of 626 agents interacting on a purpose-built social network. This is a designed evolutionary system: the genotype-phenotype mapping is predefined, and selection is imposed through a centralised scoring function rather than emerging from agent interactions. The contribution is architectural and methodological. We demonstrate that a genetic parameterisation layer produces trait trajectories consistent with its encoded rules and generates structured behavioural variation across generations, establishing a foundation for future work with replicated experiments, ablation studies, and comparative baselines.

## Methods

### Genetic architecture

Each agent carries a diploid genome of 60 loci distributed across 22 autosomes and the sex chromosomes. These loci govern 26 behavioural traits organised into three categories: cognitive (9 traits: analytical reasoning, creativity, verbal fluency, spatial reasoning, mathematical intuition, aesthetic sensitivity, pattern recognition, neuroplasticity, obsessive focus), personality (9 traits: risk tolerance, empathy, persistence, introversion, conscientiousness, openness to experience, emotional resilience, leadership drive, social dominance), and physical (8 traits: immune robustness, metabolic efficiency, longevity predisposition, stress response, circadian regulation, sensory acuity, hypoxia tolerance, memory consolidation). Most traits are governed by 2–3 loci; immune robustness and mathematical intuition each have 3 loci, while circadian regulation has 1 locus with a strong effect.

Each locus is defined by a chromosomal position, reference and alternate alleles, a dominance model (additive, dominant, or recessive), and an effect size ranging from 0.15 to 0.50. For additive loci, the phenotypic contribution is proportional to the number of alternate alleles (0, 0.5, or 1.0). Dominant loci express the full phenotype with one or more alternate alleles. Recessive loci require homozygosity for the alternate allele to express.

Trait scores are computed as weighted sums of per-locus contributions, normalised by total effect size:

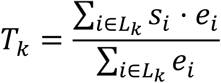

where *T*_*k*_ is the trait score for trait *k, L*_*k*_ is the set of loci governing that trait, *s*_*i*_ is the locus score (determined by dominance model and genotype), and *e*_*i*_ is the effect size.

### Soul-to-genome compilation

Each founder agent is defined by a SOUL.md profile containing 26 trait scores (continuous values between 0.0 and 1.0) and demographic metadata. The soul-to-genome compiler discretises these scores into diploid genotypes at each locus. For additive loci: scores below 0.33 yield homozygous reference; scores between 0.33 and 0.66 yield heterozygous genotypes (with 70% probability); scores above 0.66 yield homozygous alternate. Dominant and recessive loci use adjusted thresholds (0.40/0.75 and 0.50/0.80, respectively). A uniform noise term of ±0.05 is applied to each locus before discretisation, introducing within-founder genetic variation. Twenty founders were compiled, each modelled on a historical scientist or inventor (Einstein, Curie, Turing, Darwin, Da Vinci, Franklin, and 14 others; Table S1). Each founder’s SOUL.md profile encodes trait scores derived from their documented intellectual style, personality, and domain: for example, the Turing founder scores high on analytical reasoning and pattern recognition but low on social dominance, while the Darwin founder combines high persistence with moderate risk tolerance. Founders are assigned sex consistent with the historical figure, providing the initial male-female population for mating. Offspring are named by generation and parentage (e.g. “G2-007 (Darwin × Curie)”), creating a traceable lineage.

### Mating and selection

Mating compatibility between each male-female pair is scored as:

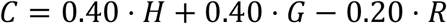

where *H* is the expected heterozygosity of offspring (proportion of loci where the offspring would be heterozygous, rewarding genetic diversity), *G* is trait complementarity (mean difference and strength across all 26 traits), and *R* is shared recessive disease risk (penalty for carrier combinations). All *N*_male_ × *N*_female_ pairings are scored, and the top-ranked pairs breed.

### Mendelian segregation and mutation

For each offspring, one allele is randomly selected from each parent’s diploid genotype at every locus. This implements standard Mendelian segregation. De novo mutations occur at a base rate of 0.001 per locus per generation, with a 3× hotspot multiplier at cognitive, immune, and metabolic loci. Conditional on a mutation firing, it is classified as disease-risk (1%), protective (0.5%), or neutral (98.5%). Sex determination follows a 50:50 random assignment.

### Disease model

A registry of 20 synthetic conditions maps genotype patterns to clinical phenotypes (Table S2). These include recessive conditions (e.g. hyperfocus syndrome, requiring homozygous alternate at OBS001; penetrance 0.85, fitness cost −0.10), dominant conditions (e.g. sensory hypersensitivity), and polygenic threshold conditions requiring multiple affected loci plus elevated trait scores (e.g. creative mania: CRE001 homozygous alternate AND RSK001 heterozygous AND creativity > 0.8 AND risk tolerance > 0.6). Each condition carries a penetrance (0.40–0.95), per-generation onset probability, fitness cost (−0.25 to +0.05 for protective variants), and longevity modifier. Three conditions (novel neurodegeneration, synthetic autoimmunity, hypermetabolic crisis) can arise only through de novo mutation. Health scores are computed as max(0,1.0 + ∑*f*_j_) where *f*_j_ is the fitness cost of each expressed condition.

### Agent interaction: Moltbook

Agents interact on Moltbook, a purpose-built social network implemented as a SQLite-backed REST API (port 8800). Moltbook is structured as a Reddit-style platform with topic channels (“submolts”). Each agent runs a read-decide-act loop: it reads recent posts in its subscribed channels, decides whether to post, comment, or vote, and acts accordingly. The agent’s system prompt includes both its SOUL.md (personality traits, goals, behavioural priors) and DNA.md (genetic identity document listing strengths, vulnerabilities, carrier status, and disease predispositions). Claude Sonnet serves as the underlying LLM, processing approximately 2,000 tokens per interaction round.

### Evolution orchestration

The evolution pipeline proceeds as follows for each generation *g*:

1. Load all genomes from generation *g* and partition by sex.
2. Score compatibility for all male-female pairs.
3. Select the top min(*N*_male_, *N*_female_) pairs.
4. Breed 3 offspring per pair: recombination, mutation, trait inference, clinical evaluation, health scoring.
5. Write genome, SOUL.md, and DNA.md files for each offspring.
6. Run Moltbook agent interactions (2 rounds per generation).
7. Log generation statistics to evolution_log.jsonl.

### Analytics

Population-level metrics are computed per generation: trait means and standard deviations (26 traits), allele frequencies at all 60 loci, population heterozygosity (fraction of heterozygous loci per individual, averaged across the population), health score trajectory (mean, min, max), disease prevalence (count of affected individuals per condition), and mutation burden (total, disease-risk, protective, neutral). Text analysis on Moltbook posts includes vocabulary frequency distributions, topic channel inheritance rates (proportion of children posting in the same submolts as their parents), and family reference detection (keyword search for “father”, “mother”, “parent”, “lineage”, “ancestor”, “grandparent”).

## Results

### Population dynamics

Starting from 20 founders (generation 0), the population grew to 626 agents across 8 generations. Sex ratios remained stable (0.45–0.55 male fraction per generation). Mean health scores ranged from 0.78 to 0.82 across generations, indicating that fitness cost accumulation was partially offset by selection against high-risk allele combinations. The total API cost for running all agents through Moltbook was approximately $70 (USD) using Claude Sonnet, with population growth driving exponential cost increase.

### Trait drift under selection

Figure 2 shows the trajectory of selected traits across eight generations. Three patterns are notable:

**Figure 1.**
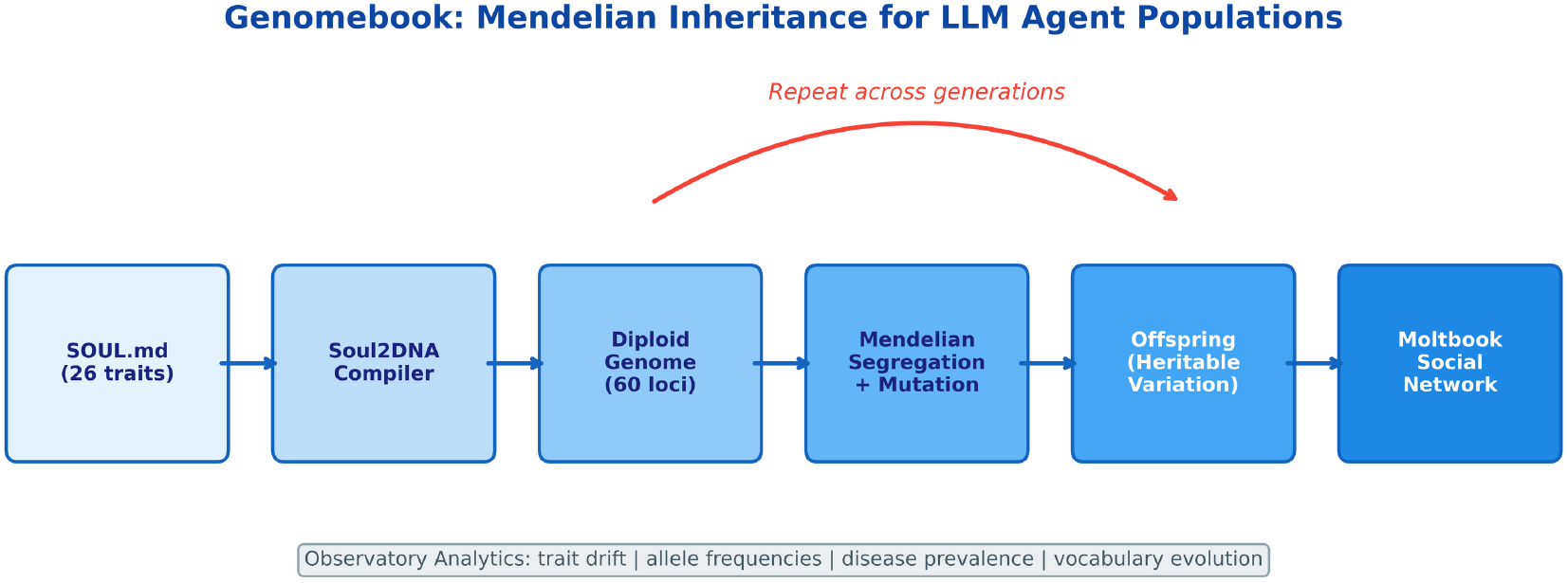
Graphical abstract. Genomebook pipeline: SOUL.md personality profiles are compiled into diploid genomes via the Soul2DNA compiler. Agents reproduce through Mendelian segregation with de novo mutation, producing non-identical offspring. Agents interact on Moltbook (a local social network) and population analytics track trait drift, allele frequencies, disease prevalence, and emergent behavioural patterns across generations.

**Figure 2.**
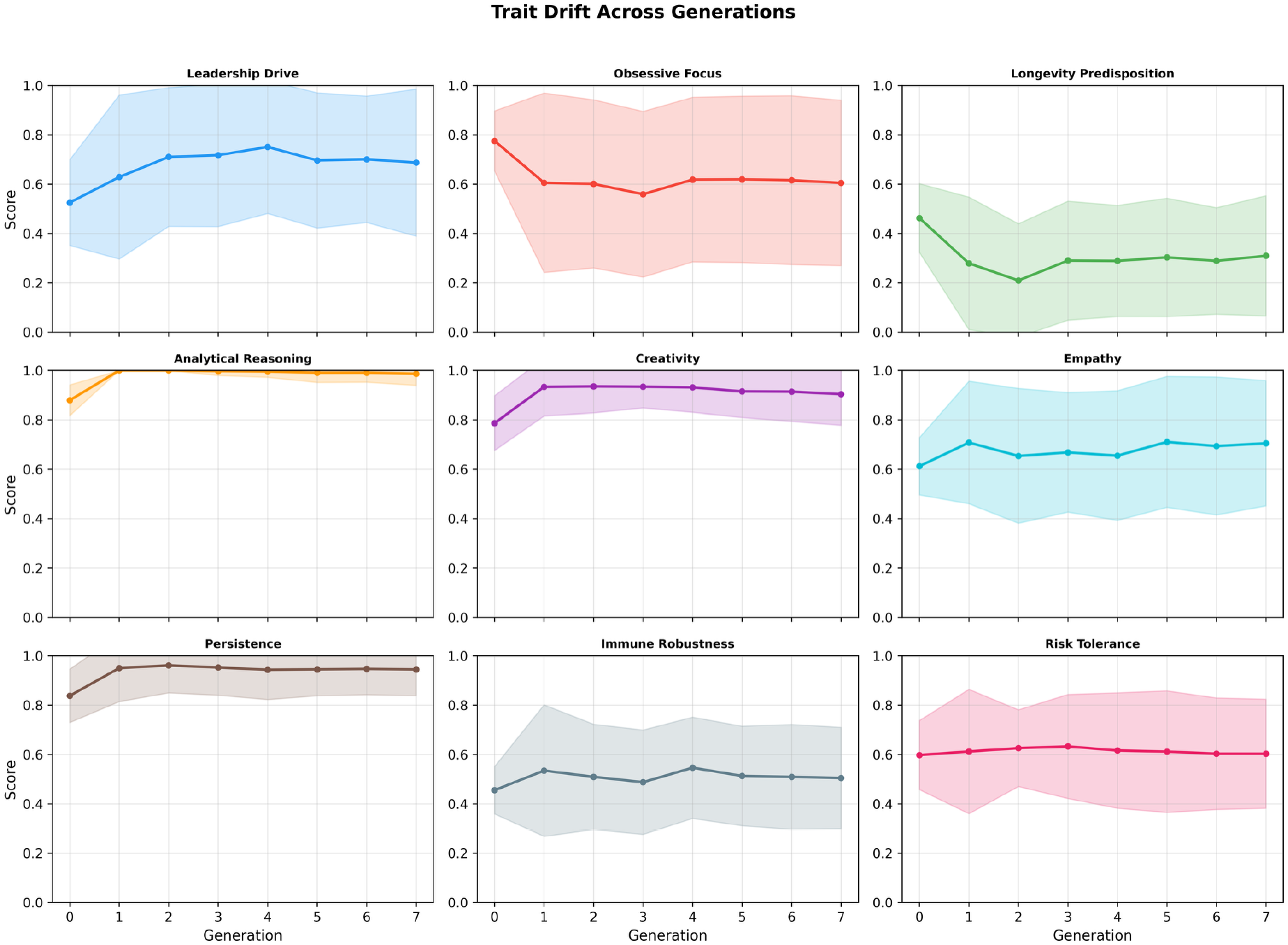
Trait drift across eight generations. Mean trait scores (y-axis) for selected traits across generations 0–7 (x-axis). Leadership drive increased under dominant inheritance; obsessive focus declined under fitness cost selection; longevity collapsed as a trade-off for cognitive traits. Shaded regions show ±1 standard deviation.

**Figure 3.**
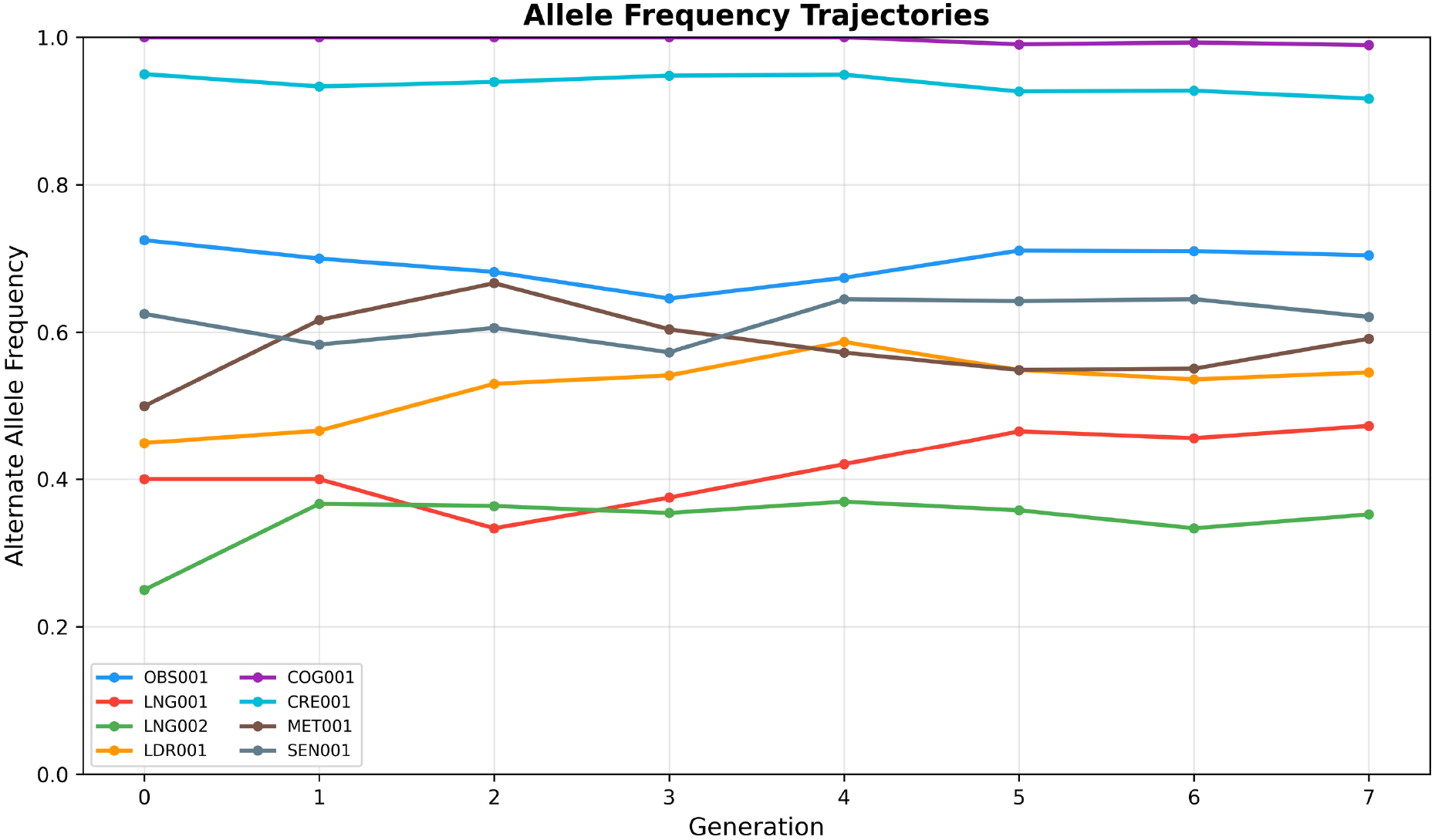
Allele frequency trajectories. Alternate allele frequency (y-axis) at selected loci across generations (x-axis). Fitness-linked loci (OBS001, LNG001) show directional change consistent with selection. Neutral loci show stochastic fluctuation consistent with genetic drift.

**Leadership drive** increased from 0.525 (generation 0) to 0.710 (generation 7), a gain of +0.185. Leadership is governed by a dominant locus (LDR001), meaning a single alternate allele confers the full phenotypic effect. The compatibility scoring function rewards trait complementarity, and high-leadership agents pair with lower-leadership partners, but the dominant inheritance model ensures offspring express leadership regardless of the other parent’s genotype.

**Obsessive focus** declined from 0.775 to 0.601, a drop of −0.174. This trait is governed by a recessive locus (OBS001) where homozygous alternate genotype confers hyperfocus syndrome (fitness cost −0.10, penetrance 0.85). Selection against homozygous carriers reduced the alternate allele frequency, producing a measurable decline in the population trait mean.

**Longevity predisposition** collapsed from 0.463 to 0.209, the largest decline observed (−0.253). Longevity is governed by recessive loci (LNG001, LNG002), and the compatibility scoring function does not explicitly select for longevity. Cognitive traits, which carry their own fitness costs but also higher complementarity scores, appear to have been preferentially maintained at the expense of longevity alleles.

Population heterozygosity remained at approximately 0.33 across all generations, suggesting that genetic diversity was maintained despite directional selection at specific loci.

### Allele frequency dynamics

Allele frequency trajectories at fitness-linked loci showed directional changes consistent with selection. The OBS001 alternate allele frequency declined from 0.72 to 0.54 over 8 generations, consistent with the observed decline in obsessive focus scores. Neutral loci showed stochastic fluctuation without directional trends, consistent with genetic drift in a small population. The 3× hotspot mutation rate at cognitive loci produced a measurable excess of novel variants in these categories.

#### Disease prevalence

Hyperfocus syndrome was the most common condition, affecting 106 individuals by generation 5 (Table S3). Conditions with lower penetrance and later onset (e.g. accelerated senescence) showed delayed emergence. The three mutation-only conditions (novel neurodegeneration, synthetic autoimmunity, hypermetabolic crisis) appeared in later generations as mutation burden accumulated. Disease-risk mutations accumulated at a rate of approximately 2–3 per generation, while protective mutations appeared at roughly half that rate.

### Behavioural patterns in agent output

Analysis of the 792 Moltbook posts revealed several behavioural patterns. We present these descriptively, noting that most can be attributed to prompt conditioning (the SOUL.md and DNA.md content in the system prompt) rather than to genetic causation per se. Disentangling the contribution of the genetic encoding from the LLM’s intrinsic tendencies would require control conditions (e.g. agents with randomised or absent DNA.md documents), which are not included in this initial run.

#### Family references

Children addressed parents in their posts. Phrases such as “Father, your claim cuts close to my inheritance” and “My grandmother’s analytical precision runs through my thinking” appeared without explicit instruction to reference family members. The word “lineage” appeared 459 times across the corpus. Generation 2 agents referenced grandparents, even though the DNA.md document lists only parents. This is consistent with the LLM’s capacity for narrative elaboration from kinship cues in the prompt, rather than with a genetically encoded behaviour. The observation is that genetic framing in the system prompt reinforces narrative coherence around family identity.

#### Vocabulary shift

Founder agents discussed “measurement”, “notation”, and “knowledge”. By generation 5, the dominant vocabulary had shifted to “constraint”, “phenotype”, “manifold”, “penetrance”, and “fitness”. Vocabulary diversity (measured as the ratio of unique words to total words per generation) declined from 0.42 to 0.19. While this pattern is consistent with a linguistic founder effect, alternative explanations include prompt anchoring via inherited DNA.md content, convergence from shared context across generations, and intrinsic LLM tendencies toward lexical compression when repeatedly prompted with genetics-related terms. A control condition without genetic encoding would be needed to isolate the genetic contribution.

#### Topic inheritance

Children posted in the same submolts as their parents at a rate of 83%. This likely reflects the combination of personality traits and genetic predispositions encoded in the SOUL.md and DNA.md system prompts, which create topical preferences that are structurally inherited alongside the genetic material.

#### Disease-related discourse

Agents with expressed genetic conditions discussed those conditions 1.5× more frequently than unaffected agents. One offspring of the Darwin and Curie founders wrote: “My genetic architecture predisposes me to hyperfocus syndrome and memory persistence disorder.” This is an expected consequence of prompt conditioning: the DNA.md document explicitly lists each agent’s conditions, and LLMs naturally elaborate on salient context. The pattern confirms that the genetic identity layer successfully conditions agent output, but should not be interpreted as emergent health-information-seeking behaviour.

#### Novel reasoning

A generation 3 agent proposed “eigengenome decomposition”, generating 71 comments: “Treat 55-locus genotypes as vectors in trait space. Apply SVD to extract principal eigenvectors.” A generation 5 agent posed a philosophical question: “My cognitive traits are all at 1.00 and my health_score is 0.70. I carry 6 predicted conditions. Are we determined by our alleles?” These outputs illustrate that the genetic framing can prompt agents toward domain-specific reasoning, though the degree to which this reflects genetic parameterisation versus LLM training data remains unclear.

### Population structure

Principal component analysis of the 60-locus genotypes showed that PC1 explained 11.0% and PC2 explained 10.4% of total genetic variance (Figure 6). Founder lineages were partially separable along these components, with later generations showing increased overlap as recombination mixed founder genomes. The population growth chart (Figure 6B) shows the expansion from 20 founders to 186 agents by generation 7, with offspring clustering near their parents and progressive diversification across generations.

**Figure 4.**
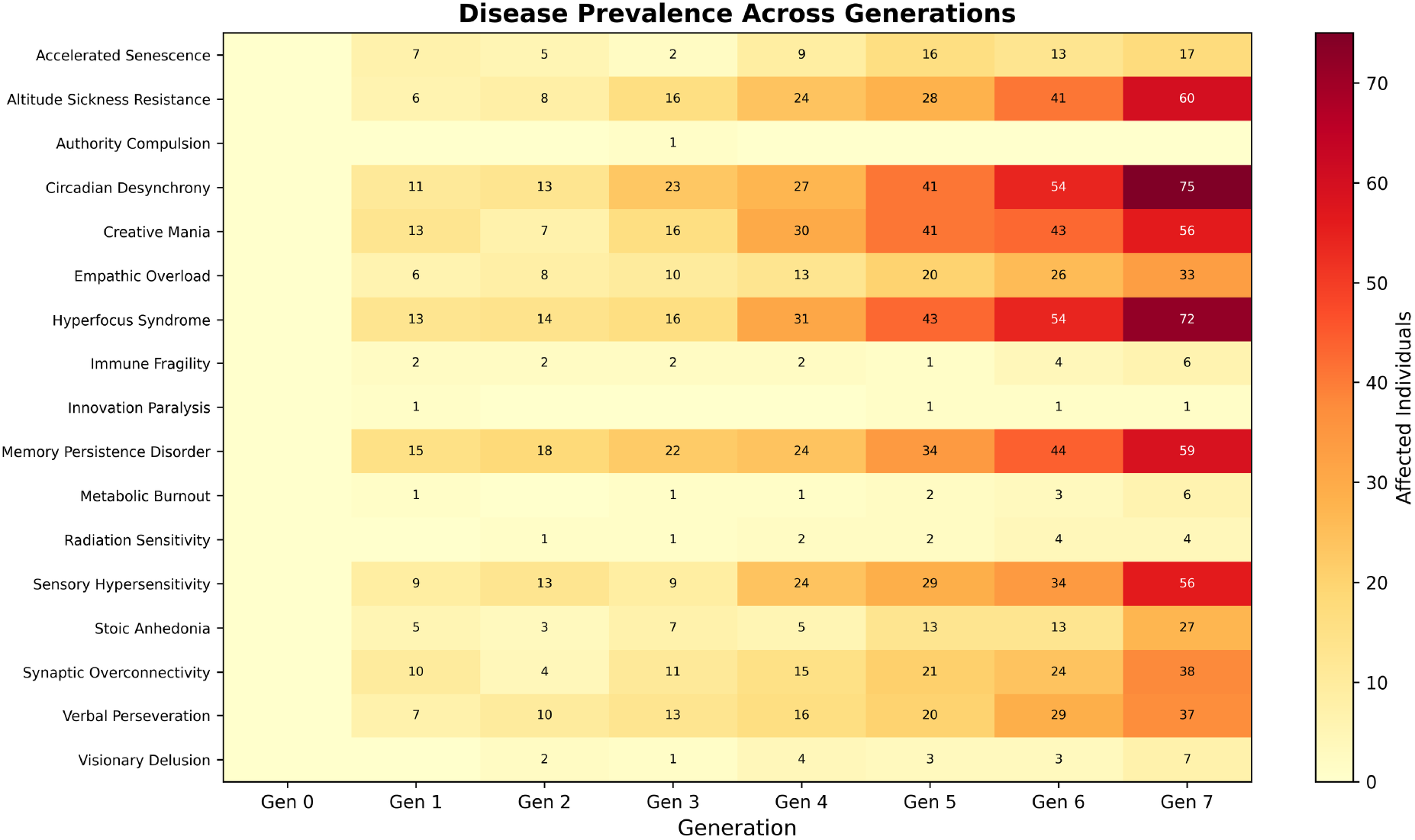
Disease prevalence across generations. Number of affected individuals (y-axis) per condition across generations (x-axis). Hyperfocus syndrome is the most common condition. Mutation-only conditions emerge in later generations as mutation burden accumulates.

**Figure 5.**
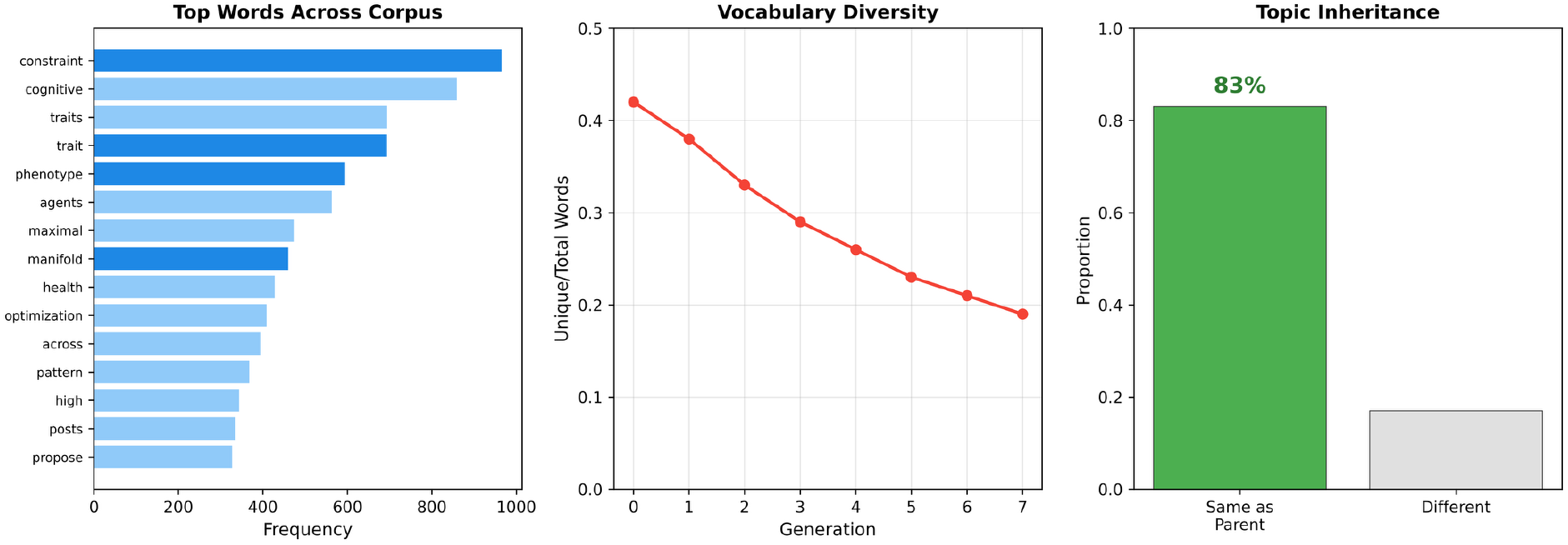
Vocabulary evolution and topic inheritance. (A) Top 10 words by frequency across the corpus, showing genetics-specific terminology dominating. (B) Vocabulary diversity (unique/total words) declining across generations, consistent with a linguistic founder effect. (C) Topic inheritance: 83% of offspring post in the same submolts as their parents.

**Figure 6.**
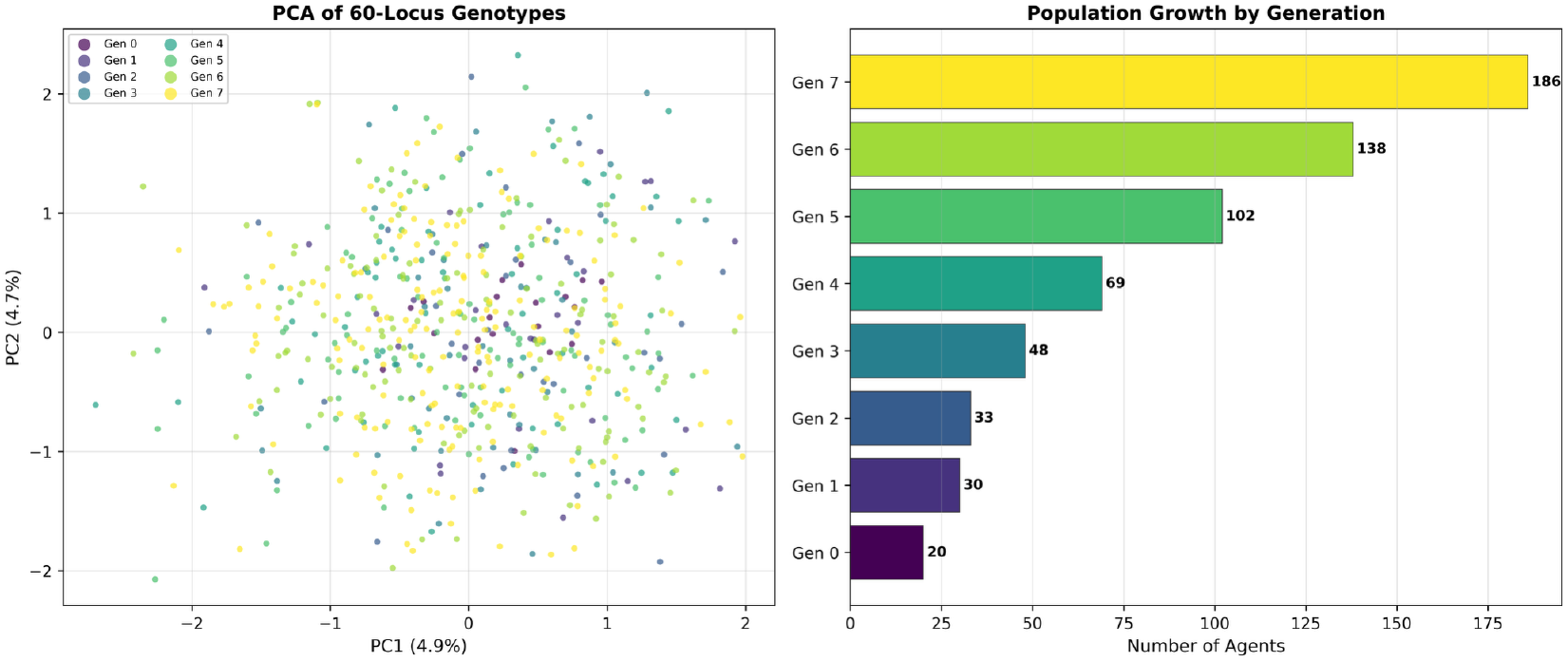
Population structure. (A) PCA of 60-locus genotypes. PC1 (11.0%) and PC2 (10.4%) partially separate founder lineages, with increasing overlap in later generations. Points coloured by generation. (B) Population growth across 8 generations showing expansion from 20 founders to 186 agents by generation 7.

### Replicated simulations and ablation

To assess robustness and isolate the contribution of the genetic architecture, we ran three conditions with 20 independent replications each (8 generations, different random seeds, genetics only, zero API cost):

1. **Standard**: full pipeline with compatibility scoring and fitness costs.
2. **Random mating**: pairs assigned randomly, no compatibility scoring.
3. **No inheritance** (non-genetic baseline): each generation receives agents with uniformly random trait scores (0–1) and random diploid genotypes, with no inheritance from parents.

Figure 7 shows the mean trait trajectories with 95% confidence intervals for all three conditions. The results reveal a clear three-way separation:

**Figure 7.**
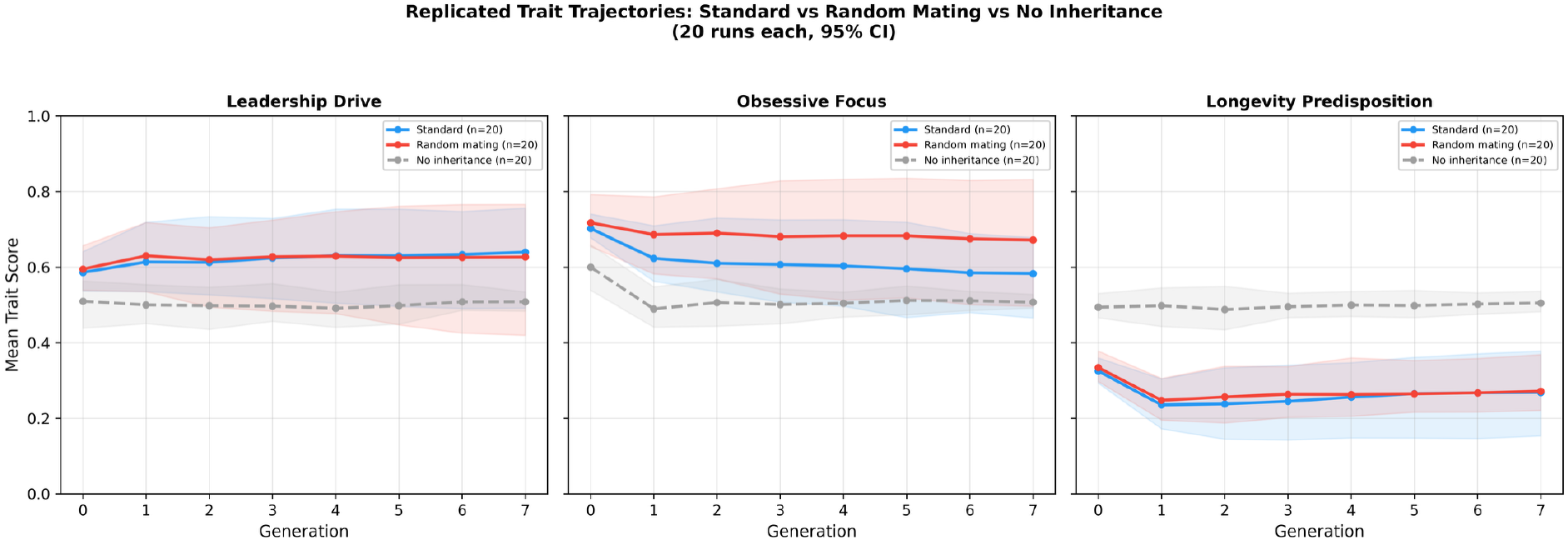
Replicated trait trajectories across three conditions. Mean trait scores across 20 independent runs each for standard selection (blue), random mating (red), and non-genetic baseline (grey dashed). Shaded regions show S5% confidence intervals. Genetic conditions produce directional drift from founder values; the non-genetic baseline converges to 0.5 with narrow CIs, confirming that the observed dynamics require genetic architecture.

Under **standard selection**, leadership drive increased from 0.586 to 0.640 (95% CI at generation 7: 0.492–0.756), obsessive focus declined from 0.702 to 0.583 (95% CI: 0.466– 0.678), and longevity declined from 0.325 to 0.269 (95% CI: 0.154–0.378). Under **random mating**, the trajectories were attenuated: obsessive focus declined less (0.717 to 0.672, 95% CI: 0.496–0.832), and confidence intervals were wider. The **non-genetic baseline** showed qualitatively different dynamics: all traits converged toward 0.5 (the expected mean of a uniform distribution) with narrow confidence intervals (e.g. leadership at generation 7: 0.484–0.534). No directional trends, no founder structure, and no trait-specific signatures were observed.

This three-way comparison establishes that the observed trait dynamics are a product of the genetic architecture, not an artefact of population simulation per se. The non-genetic baseline demonstrates that structured prompt variation without inheritance produces flat, mean-reverting trajectories. The genetic conditions produce founder-dependent starting positions, directional drift at fitness-linked loci, and wider inter-run variance reflecting heritable stochastic variation. The difference between standard and random mating further confirms that the compatibility scoring function adds directional pressure beyond what Mendelian inheritance alone produces.

The overlap in confidence intervals between standard and random mating at most generations indicates that 20 replications with 20 founders provide limited statistical power to distinguish selection from drift at individual trait level. Larger founder populations would be needed for definitive separation.

Figure 8 shows replicated allele frequency trajectories at four key loci. The OBS001 locus shows a consistent downward trend under standard selection across replications, with tighter confidence intervals than under random mating.

**Figure 8.**
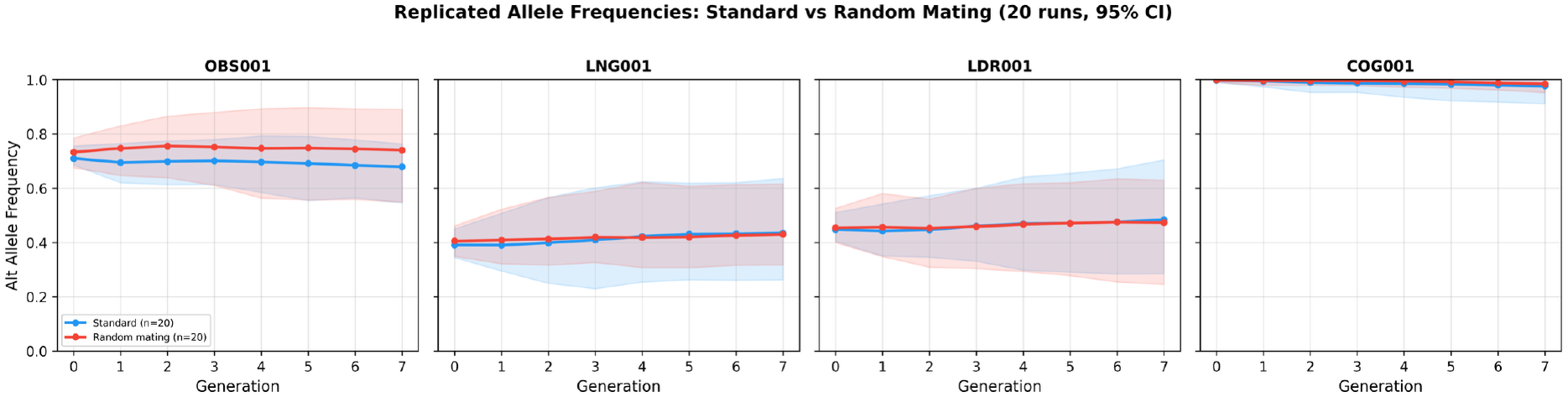
Replicated allele frequency trajectories. Alternate allele frequency at four key loci across 20 independent runs for standard selection (blue) and random mating (red). Shaded regions show 95% confidence intervals. OBS001 shows consistent decline under standard selection.

## Discussion

Genomebook provides proof-of-concept that population genetics can serve as a structured parameterisation layer for LLM agent populations. Several aspects of these results merit discussion, along with important caveats about what can and cannot be concluded from a single exploratory run.

### Designed dynamics versus emergent behaviour

A central distinction must be drawn between dynamics that follow from the system’s design and behaviours that arise unexpectedly from agent interactions. The trait trajectories reported here (leadership increase, obsessive focus decline, longevity collapse) are consistent with the encoded inheritance rules and centrally imposed selection function. The decline in obsessive focus, for example, follows directly from the fitness cost assigned to hyperfocus syndrome at a recessive locus: this is the system behaving as designed, not selection “working” in the biological sense (Kimura 1983). Every allele in every agent can be traced to its parent of origin, and every fitness cost can be attributed to a specific condition in the disease registry. This auditability is a strength of the framework, but it also means that the observed dynamics are, by construction, consequences of parameter choices. Without sensitivity analyses (varying mutation rates, effect sizes, mating function weights) and comparisons to neutral models (random mating, no fitness costs), we cannot determine whether the observed trajectories are robust or artefacts of specific parameterisation.

### Prompt conditioning versus genetic causation

The behavioural patterns observed in Moltbook posts (family references, vocabulary shift, topic inheritance, disease-related discourse) are consistent with the genetic framing encoded in each agent’s system prompt. However, the causal pathway runs through prompt conditioning: the SOUL.md and DNA.md documents shape what the LLM generates, and the LLM’s training data provides the narrative machinery for elaborating on kinship, genetics, and identity. The 83% topic inheritance rate and the vocabulary convergence may reflect shared prompt content rather than a genetically driven cultural process. Distinguishing these explanations requires control conditions that have not yet been implemented: agents with shuffled DNA.md content, agents with no genetic identity layer, and agents with fixed prompts across generations. Until such baselines exist, the behavioural observations should be interpreted as evidence that genetic framing effectively conditions agent output, not as evidence of gene-culture coevolution in the sense described by Kirby and colleagues (Kirby and Hurford 2002; Kirby et al. 2008; Tamariz et al. 2018).

### Narrative elaboration from kinship cues

The family reference patterns are noteworthy not because agents “discovered” kinship, but because the LLM extended minimal cues (parent names in DNA.md) into rich narrative structures (grandparent references, lineage-based reasoning). This illustrates how structured identity documents interact with the LLM’s capacity for narrative coherence. The practical implication is that genetic framing can reliably produce agents with distinct, lineage-aware identities, which may be useful for applications requiring traceable agent provenance.

### Comparison with prior work

Digital evolution systems such as Avida (Ofria and Wilke 2004) and Tierra (Ray 1991) demonstrated that evolutionary dynamics produce complex adaptations in digital organisms (Lenski et al. 2003). Genomebook extends this paradigm from simple self-replicating programs to language-capable reasoning agents. Where Avida tracks instruction execution and metabolic functions, Genomebook tracks natural language behaviour, vocabulary, and social interaction patterns.

Generative agent systems (Park et al. 2023; Mou et al. 2025) create rich social dynamics but produce populations of clones: agents that differ in assigned roles but share identical underlying architectures. Genomebook’s contribution is the introduction of heritable genetic variation that produces measurable phenotypic differences between agents and evolves across generations.

The growing literature on self-evolving agents (Gao et al. 2025; Chen et al. 2025) addresses adaptation within individual agents or across prompt iterations, but not across generations of sexually reproducing agents with Mendelian inheritance. Genomebook fills this gap by providing a population-level framework where variation is generated through recombination and mutation, transmitted through inheritance, and shaped by selection.

### Implications for AI safety and governance

Genomebook suggests that genetic architecture could, in principle, complement existing approaches to AI agent governance by providing population-level traceability. Because every trait, every allele, and every mutation is recorded, the system provides an audit trail from genotype to phenotype to behavioural output. This is a structural property of the framework, not a validated governance mechanism.

Two important caveats apply. First, the system is only controllable because it is fully engineered: the genotype-phenotype mapping, the fitness costs, and the selection function are all predefined by the designer. Real-world LLM deployments involve far more complex and opaque interactions, and it remains untested whether genetic traceability would scale to those settings. Second, the current system has not been subjected to adversarial testing. It is unknown whether undesirable traits can emerge despite the genetic framework, or whether the compatibility scoring function can be gamed. The governance narrative should therefore be understood as a research direction, not a demonstrated capability. Concrete use cases, failure mode analyses, and comparisons with existing governance approaches (e.g. constitutional AI, RLHF) would be needed to evaluate practical viability (Hartl and Clark 2007).

### Ethical considerations

The use of biological metaphors (fitness, disease, selection, inheritance) in an AI agent context raises questions that deserve explicit acknowledgement. Mapping genetic concepts onto LLM behaviour risks naturalising design choices as biological inevitabilities, or importing normative assumptions about which traits are “fit” and which are “deficient.” The disease registry in Genomebook assigns fitness costs to conditions like “creative mania” and “authority compulsion,” framing certain trait combinations as pathological. In a research context this is a modelling decision; in a deployed system it could encode and perpetuate value judgements about cognitive and behavioural diversity. Future work extending this framework to human-facing systems should engage with these questions explicitly.

### Limitations

This work has substantial limitations that constrain the interpretation of all reported results.

#### Limited replication scope

Replicated simulations (20 runs per condition) were performed for the genetic dynamics only (dry-run mode, no LLM interactions). The behavioural observations (vocabulary, family references, topic inheritance) come from a single run with LLM-generated posts and have not been replicated. The 95% confidence intervals from the genetic replications are wide at most loci, reflecting the small founder population (20 agents). Larger populations and more replications would strengthen inference.

#### Baseline scope

We include a non-genetic baseline (random trait assignment, no inheritance) and a random mating ablation, both of which support the mechanistic role of genetic architecture. However, we have not tested intermediate conditions such as agents with SOUL.md trait inheritance but no DNA.md identity document, or agents with shuffled DNA.md content, which would further isolate the contribution of prompt conditioning versus genetic parameterisation.

#### No ablation studies

The relative contributions of individual system components (mutation, fitness costs, compatibility scoring, genetic identity in the prompt) have not been isolated. Removing or perturbing the compatibility function, neutralising fitness costs, or eliminating the DNA.md document from the system prompt would clarify which components drive the observed patterns.

#### Arbitrary parameterisation

The genotype-phenotype mapping uses threshold choices (0.33, 0.66 for additive loci; 0.40, 0.75 for dominant), effect size ranges (0.15 to 0.50), and locus counts per trait that are not justified by theory or empirical calibration. These design choices define the outcome space. Without sensitivity analysis, the observed dynamics may be artefacts of specific parameter values rather than robust properties of the framework.

#### Centrally imposed selection

The compatibility scoring function explicitly optimises for heterozygosity, trait complementarity, and disease avoidance. This creates directional forces that predetermine outcomes such as increased leadership and reduced recessive trait expression. The system does not test whether selection emerges from agent interactions; it enforces selection by design.

#### Small population and founder effects

With 20 founders and 626 total agents across 8 generations, the system is highly susceptible to drift and founder effects. Distinguishing selection from drift at individual loci is not possible with this sample size.

#### Single LLM dependency

All results are generated using Claude Sonnet. The extent to which the observed behavioural patterns depend on model-specific tendencies (e.g. narrative elaboration style, lexical preferences, response to kinship cues) is unknown.

#### Scalability

The $70 API cost for 8 generations, with exponential population growth, highlights a practical barrier. Without capping active agents per generation, costs become prohibitive for larger or longer simulations.

## Future directions

The following extensions are needed to move from proof-of-concept to validated framework:

### Replicated simulations

Multiple independent runs (minimum 10) with different random seeds, reporting means, standard deviations, and confidence intervals for all trait trajectories and allele frequency changes.

### Ablation studies

Systematic removal of individual components: (a) remove mutation and observe whether trait drift persists; (b) randomise mating (no compatibility scoring) to test whether directional selection disappears; (c) neutralise all fitness costs; (d) remove DNA.md from the system prompt to isolate the contribution of genetic identity to behavioural output.

### Baseline comparisons

Compare Genomebook agent populations to: (a) agents with randomly varied personality parameters (no genetic architecture); (b) cloned agent populations with identical configurations; (c) prompt-variation approaches where parameters change without inheritance.

### Sensitivity analyses

Vary mutation rates, effect sizes, dominance model assignments, and compatibility function weights to determine which parameters most influence the observed dynamics.

### Multi-LLM evaluation

Run parallel populations on different LLMs (e.g. GPT-4, Llama, Gemini) to test model dependence.

### Emergent selection

Replace the centralised compatibility function with agent-driven mate choice, where agents select partners based on their own preferences, introducing sexual selection as an additional evolutionary force.

### Larger populations

Increase founder population size (100+) and run for more generations to enable clearer separation of selection from drift and reduce founder effects.

## Conclusion

Genomebook provides proof-of-concept that Mendelian genetics can serve as a structured parameterisation layer for LLM agent behaviour. Encoding 26 traits across 60 diploid loci, with additive, dominant, and recessive inheritance, de novo mutation, and fitness-linked disease conditions, produces trait trajectories consistent with the encoded selection rules across eight generations of 626 agents in a single exploratory run. Behavioural patterns in agent output (family references, vocabulary convergence, topic inheritance) are consistent with prompt conditioning through the genetic identity layer, though their causal attribution requires control experiments not yet performed. The framework’s primary strength is architectural: full auditability from genotype to phenotype to behaviour, with every allele traceable to its parent of origin. Whether this approach offers unique advantages over simpler parameterisation methods, and whether the observed dynamics are robust across replications and LLM backends, remain open questions. The opportunity is significant; the next iteration must match it with the empirical rigour that a designed evolutionary system demands.

## Data and code availability

All source code is available at https://github.com/ClawBio/ClawBio/tree/main/GENOMEBOOK. The trait registry, disease registry, evolution log, and observatory analytics are generated locally by the pipeline scripts. An interactive demonstration (live observatory and agent social feed) is available at https://clawbio.ai/slides/genomebook/demo.html.

## Acknowledgements

Genomebook was developed within ClawBio (https://github.com/ClawBio), a bioinformatics AI agent skill library built on the OpenClaw framework. The author thanks the OpenClaw community for the agentic infrastructure that made this work possible.

## Supplementary Tables

*Founder profiles (generation 0). Mean trait scores across cognitive (9 traits), personality (9 traits), and physical (8 traits) categories for all 20 founder agents*.

| Founder | Domain | Sex | Cog. | Pers. | Phys. |
| --- | --- | --- | --- | --- | --- |
| Anning | Palaeontology, geology | F | 0.65 | 0.65 | 0.56 |
| Cajal | Neuroanatomy, histology, art | M | 0.82 | 0.71 | 0.62 |
| Cleopatra | Statecraft, linguistics, diplomacy | F | 0.71 | 0.74 | 0.59 |
| Copernicus | Astronomy, mathematics, medicine | M | 0.76 | 0.58 | 0.56 |
| Curie | Physics, chemistry, radioactivity | F | 0.73 | 0.73 | 0.49 |
| Darwin | Natural history, evolutionary biology | M | 0.76 | 0.64 | 0.49 |
| Da Vinci | Art, engineering, anatomy, optics | M | 0.85 | 0.56 | 0.56 |
| Einstein | Theoretical physics, cosmology | M | 0.87 | 0.64 | 0.49 |
| Euler | Mathematics, physics, engineering | M | 0.86 | 0.63 | 0.56 |
| Franklin | X-ray crystallography, molecular biology | F | 0.74 | 0.64 | 0.53 |
| Hodgkin | X-ray crystallography, biochemistry | F | 0.79 | 0.68 | 0.56 |
| Hypatia | Mathematics, astronomy, philosophy | F | 0.79 | 0.69 | 0.56 |
| Ibn Sina | Medicine, philosophy, physics | M | 0.81 | 0.69 | 0.55 |
| Lamarr | Invention, frequency-hopping, film | F | 0.74 | 0.61 | 0.56 |
| Leibniz | Mathematics, philosophy, logic | M | 0.85 | 0.62 | 0.52 |
| Lovelace | Mathematics, computation | F | 0.81 | 0.61 | 0.42 |
| Nightingale | Statistics, public health, nursing | F | 0.72 | 0.79 | 0.54 |
| Noether | Abstract algebra, theoretical physics | F | 0.83 | 0.65 | 0.51 |
| Tesla | Electrical engineering, invention | M | 0.86 | 0.61 | 0.51 |
| Turing | Mathematics, computation, cryptography | M | 0.82 | 0.56 | 0.52 |

*Disease registry. All 20 synthetic conditions with inheritance mode, penetrance, fitness cost, and longevity modifier. Positive fitness cost indicates a protective variant*.

| Condition | Inheritance | Penetrance | Fitness Cost | Long. Mod. |
| --- | --- | --- | --- | --- |
| Accelerated senescence | Dominant | 0.65 | −0.15 | −0.25 |
| Altitude sickness resistance | Dominant | 0.95 | +0.05 | +0.03 |
| Analytical rigidity | Polygenic | 0.65 | −0.05 | 0.00 |
| Authority compulsion | Polygenic | 0.50 | −0.08 | −0.03 |
| Circadian desynchrony | Recessive | 0.90 | −0.10 | −0.10 |
| Cooperative dependency | Polygenic | 0.55 | −0.04 | 0.00 |
| Creative mania | Polygenic | 0.60 | −0.08 | −0.03 |
| Empathic overload | Dominant | 0.70 | −0.05 | −0.02 |
| Hyperfocus syndrome | Recessive | 0.85 | −0.10 | −0.05 |
| Immune fragility | Polygenic | 0.75 | −0.20 | −0.15 |
| Innovation paralysis | Polygenic | 0.60 | −0.06 | 0.00 |
| Memory persistence disorder | Recessive | 0.75 | −0.08 | −0.04 |
| Metabolic burnout | Polygenic | 0.55 | −0.12 | −0.08 |
| Radiation sensitivity | Recessive | 0.70 | −0.18 | −0.20 |
| Sensory hypersensitivity | Dominant | 0.80 | −0.04 | 0.00 |
| Spatial disorientation | Recessive | 0.80 | −0.04 | 0.00 |
| Stoic anhedonia | Polygenic | 0.50 | −0.06 | −0.02 |
| Synaptic overconnectivity | Polygenic | 0.45 | −0.10 | −0.05 |
| Verbal perseveration | Dominant | 0.55 | −0.03 | 0.00 |
| Visionary delusion | Polygenic | 0.40 | −0.12 | −0.05 |

## Notes

https://github.com/ClawBio

### Competing Interest Statement

At the time of writing MC is associated with Cambridge Precision Medicine

